# The membrane skeleton density of red blood cells in *MYH9*-related disease patients is decreased

**DOI:** 10.1101/2025.02.11.637639

**Authors:** Shaopeng Sun, Jiabin Pan, Ning Zhang, Xianghong Jin, Tie-nan Zhu, Zhijie Kang, Jihong Hao, Xiang-dong Li

**Affiliations:** Group of Cell Motility and Muscle Contraction, State Key Laboratory of Integrated Management of Pest Insects and Rodents, Institute of Zoology, Chinese Academy of Sciences, Beijing 100101, China; University of Chinese Academy of Sciences, Beijing 100049, China; Department of Rare Diseases, Peking Union Medical College Hospital, Chinese Academy of Medical Science & Peking Union Medical College, Beijing, China; Department of Hematology, Peking Union Medical College Hospital, Chinese Academy of Medical Sciences & Peking Union Medical College, Beijing 100730, China; Department of Hematology, the Second Hospital of Dalian Medical University; Liaoning Medical Center for Hematopoietic Stem Cell Transplantation; Liaoning Key Laboratory of Hematopoietic Stem Cell Transplantation and Translational Medicine, Dalian, China; Department of Clinical Laboratory, the Second Hospital of Hebei Medical University, Shijiazhuang 050000, China

## Abstract

*MYH9*-related disease (*MYH9*-RD) is a rare autosomal dominant disorder caused by mutations in *MYH9* gene, which encodes the heavy chain of nonmuscle myosin IIA. Nearly all *MYH9*-RD patients present with macrothrombocytopenia, characterized by decreased platelet count and increased platelet size. In this study, we collected blood samples from three *MYH9*-RD patients (R702S, D1424N, and R1464C) and unexpectedly found that the actin levels in the red blood cells (RBCs) from all three *MYH9*-RD patients are substantially lower than the healthy controls. We further revealed that the levels of two RBC membrane skeleton proteins, α-spectrin and tropomodulin, are also reduced in *MYH9*-RD RBCs. We showed that the membrane skeleton of *MYH9*-RD RBCs was more porous and that *MYH*9-RD RBCs produced more severe deformation under hyperosmotic pressure compared to healthy controls. We conclude that *MYH9*-RD mutations reduce the RBC membrane skeleton density and impair its mechanical properties, and propose that defects in the membrane skeleton network in RBCs may be a common symptom in *MYH9*-RD.

**Key points:** The levels of membrane skeleton proteins and the density of membrane skeleton network in RBCs of *MYH9*-RD patients are reduced.

*MYH9*-RD RBCs produce greater deformation under hypertonic conditions compared to healthy controls.

## Introduction

*MYH9*-related disease (*MYH9*-RD) is a rare autosomal dominant disorder caused by mutations in *MYH9* gene ^1-3^. Although different *MYH9*-RD mutations have different effects on disease severity and progression, almost all *MYH9*-RD patients develop macrothrombocytopenia, which is characterized by a decrease in platelet count and an increase in platelet size ^4^. Additionally, some patients may later develop non-syndromic deafness, nephritis, and cataracts ^1,5,6^. A recent study reported that *MYH9* mutations can also lead to abnormalities in red blood cell (RBC) morphology ^7^.

The *MYH9* gene encodes the heavy chain (HC) of nonmuscle myosin IIA (NMIIA), an actin-based hexameric motor protein composed of two HCs, two essential light chains (ELCs), and two regulatory light chains (RLCs) ^8^. Each HC contains a globular motor head that possesses actin binding and ATPase activities, a lever arm stabilized by one ELC and one RLC, and a C-terminal α-helical tail that dimerizes the two HCs, forming the coiled-coil tail of the hexamer. NMIIA is widely expressed and plays a crucial role in a broad range of fundamental cellular processes, including cell migration, cytokinesis, morphological changes, and adhesion ^8-10^. Over 200 pathogenic mutations associated with NMIIA have been reported, the majority of which are missense mutations. These mutations can be broadly categorized into two main types based on their location within the structural domains: mutations in the motor head and those in the tail region. The most frequent and pathogenic mutations in motor domain affect the Arg702 (R702), which are associated with the most severe and complex symptoms in *MYH9*-RD patients ^11-14^. Among the mutations in the tail region, mutations at Asp1424 (D1424) are one of the most prevalent and extensively studied tail mutations ^12,13^. While different *MYH9*-RD mutations have been reported to differentially affect the structure and function of NMIIA, it appears that many *MYH9*-RD mutations result in elevated NMIIA activity ^10,15-19^. The myosin II inhibitor blebbistatin was reported to rescue the proplatelet formation defects in *in vitro*-generated megakaryocytes from 11 patients with different *MYH9*-RD mutations ^20^.

Similar to the situation in platelets, NMIIA is the only NMII isoform present in RBCs ^7,21^. However, unlike the obvious platelet defects in *MYH9*-RD patients, the symptoms in *MYH9*-RD RBCs are very mild. *MYH9*-RD does not cause clinically significant anemia and patient RBCs have normal osmotic deformability except for a slight decrease of hemoglobin content and a slight increase in the number of elongated RBCs ^7^. RBCs are biconcave discs, and the maintenance of their shape and deformability relies on the membrane skeleton located beneath the plasma membrane ^22-24^. The membrane skeleton in RBCs is a highly cross-linked two-dimensional network composed of flexible (α1β1)_2_-spectrin tetramers, which interconnect at junctional complexes formed by short actin filaments (∼37 nm in length) and several associated proteins, including tropomodulin, P4.1, and dematin ^22,25-29^. RBC membrane skeleton is regularly organized with a constant proportion of its constituent proteins ^27,30^. Smith et al. demonstrated that NMIIA forms bipolar filaments that associate with the membrane cytoskeleton through their motor domains, generating tension via motor activities to control the biconcave disc shape and deformability of RBCs ^31^. It was proposed that *MYH9*-RD mutations enhance the association of NMIIA with spectrin-actin membrane skeleton, causing abnormal RBC morphology ^7^. However, given that the density of NMIIA filaments in the RBC is much lower than the density of spectrin-actin membrane skeleton, it is puzzling that NMIIA contractility has such a large effect on the shape and membrane properties of RBCs ^32^.

In this study, we collected blood samples from three *MYH9*-RD patients (R702S, D1424N, and R1464C) and unexpectedly found that the actin levels in the RBCs from all three *MYH9*-RD patients are substantially lower than normal subjects. We further revealed that the levels of RBC membrane skeleton proteins α-spectrin and tropomodulin are also reduced in *MYH9*-RD RBCs. We showed that the membrane skeleton of *MYH9*-RD RBCs is more porous and that *MYH9*-RD RBCs produce more severe deformation under hyperosmotic pressure compared to normal subjects. We conclude that *MYH9*-RD mutations reduce the RBC membrane skeleton density and impair its mechanical properties.

## Methods

### Blood collection

Whole blood was collected from *MYH9*-RD patients or healthy human donors into EDTA tubes at the Second Hospital of Hebei Medical University (Shijiazhuang, China), Peking Union Medical College Hospital (Beijing, China), the Second Hospital of Dalian Medical University (Dalian, China). Complete blood counts (CBCs) were determined with automated hematology analyzers at the point of collection and peripheral blood smears were prepared and stained with Wright-Giemsa. Additional EDTA tubes containing whole blood were collected at the same time and delivered within 24 hours at 4°C to Institute of Zoology, Chinese Academy of Sciences (Beijing, China), where osmotic fragility assays, RBC isolation, Wright-Giemsa-stained peripheral blood smears, and freeze-etching (see below) were performed within 24 h of arrival. All remaining blood samples were aliquoted into small volume, flash-frozen in liquid nitrogen and stored at -80°C. Blood samples were retrieved from patients upon informed consent, in accordance with the Declaration of Helsinki (approval number of Research Ethics Committee of the Second Hospital of Hebei Medical University: 2022-R030).

### SDS-PAGE and Western blot analysis of whole blood and RBCs

The whole blood samples of *MYH9*-RD patients and healthy controls for SDS-PAGE and Western blot analysis were prepared as follows. Whole blood was diluted 10 times by mixing 100 μL of whole blood with 900 μL of ice-cold lysis buffer (50 mM Tris pH 7.4, 150 mM NaCl, 1% Triton X-100, 5 mM EDTA, 10 mM NaF, 0.5% sodium deoxycholate, 0.1% SDS, 0.2 mM PMSF, 1x protease and phosphatase inhibitors). After vigorously vortexed, the diluted whole blood sample was incubated on ice for 20 minutes and then sonicated at 4°C to ensure complete lysis. The lysed sample was mixed with equal volume of 2x SDS loading buffer (100 mM Tris-HCl pH 6.8, 4% SDS, 0.08% bromophenol blue, 24% glycerol, 2% β-mercaptoethanol) and then incubated at 100°C for 5 minutes. The whole blood samples were aliquoted into small volume and stored at -80°C.

For SDS-PAGE analysis, 10 μL of whole blood samples or RBCs samples were electrophoresed on gradient SDS-polyacrylamide gels (4-20%), followed by Coomassie brilliant blue (CBB) staining. For Western blot analysis, 10 μL of blood samples were electrophoresed on SDS-polyacrylamide gels (10% gels for actin, tropomodulin, dematin and P4.1; 6% gels for α-spectrin and NMIIA HC), and then transferred to PVDF membranes for immunoblotting. Primary antibodies were diluted 1,000 times, including mouse anti-β-actin (Proteintech, 66009-1), rabbit anti-tropomodulin (Sangon Biotech, D126514), rabbit anti-dematin (Sangon Biotech, D163701), rabbit anti-P4.1 (Solarbio life science, K109011P), rabbit anti-α-spectrin (Solarbio life science, K009298P), and rabbit anti-NMIIA HC (Solarbio life science, K001583P). Secondary antibodies were diluted 10,000 times, including donkey anti-rabbit-HRP (Abcam ab205722) and donkey anti-mouse-HRP (Abcam ab6820). The protein band intensities were measured using Image J.

RBCs from *MYH9*-RD patients with D1424N or R1464C mutation were isolated immediately upon arrival. 2 mL of fresh whole blood was diluted with 2 mL of PBS, and then carefully layered over 3 mL of Ficoll-Paque PLUS (cytiva) in a centrifuge tube. The mixture was centrifuged at 400 × g for 30 minutes at 20 °C to settle the RBCs at the bottom layer. The RBCs were collected from the centrifuge tube and washed 3 times in ice-cold PBS by suspending in PBS followed by centrifugation at 600 × g for 5 min to collect the RBCs. After the final wash, the RBCs were suspended in PBS and the volume was adjusted to 2 mL, matching the initial volume (2 mL) of the whole blood samples. The isolated RBCs were aliquoted into 100 μL, quick-frozen in liquid nitrogen and stored at -80°C. The RBCs from healthy human donors were processed at the same time as a control. The RBCs samples for SDS-PAGE and Western blot were prepared following the same procedures for whole blood, except for using the isolated RBCs samples corresponding to 100 μL of whole blood.

### Quick-freeze, Deep-etch and Cryo-SEM of RBC membrane skeletons

The whole blood samples were fixed in silicon substrates immediately upon arrival as follows. Whole blood (2 μL) was diluted 100 times by gently mixing with 198 μL of PBS. The diluted whole blood (15 μL) was applied to poly-L-lysine-treated silicon substrates and incubated at room temperature for 30 minutes to facilitate RBCs adhesion. The samples were incubated with 50 mM KPO_4_, pH7.4, 2 mM MgCl_2_, 10 mM EGTA, 1 μM phallacidin, 0.5% Triton X-100, and 0.05% glutaraldehyde at room temperature for 2 min to permeabilize RBCs, and then washed rapidly with Wash buffer (50 mM KPO_4_, pH7.4, 2 mM MgCl_2_, 10 mM EGTA, 1 μM phallacidin). Subsequently, the samples were treated with 1% glutaraldehyde in wash buffer for 10 min at 37°C to fix RBC membrane skeletons. After washed with distilled water, the silicon substrates were quick frozen and stored in liquid nitrogen.

Deep-etch and replication were performed using Leica EM ACE600 Sputter Coater. The silicon substrates stored in liquid nitrogen were transferred to Leica EM ACE600 Sputter Coater using a cryopreservation transfer system (Leica EM VCT500 Vacuum Cryo Transfer System). The samples were frozen-dried at -84°C for 20 minutes and metal-shadowed with ∼4 nm of platinum. Finally, the samples were transferred to a Cryo-SEM (Zeiss crossbeam 340) for imaging under liquid nitrogen conditions. The pore size of the membrane skeleton in the central 3/4 regions of the RBCs was analyzed with Image J.

### Peripheral blood smears

The effects of different osmotic conditions on the morphology of RBCs were analyzed immediately after the arrival of blood samples. The blood samples were treated with different osmotic conditions at room temperature for 30 minutes and then stained with Wright-Giemsa. The blood smears were imaged using a Nikon ECLIPSE 80i with a 100x oil immersion objective.

### Statistical analyses

The significance of differences among means was assessed using one-way ANOVA followed by a Bonferroni post hoc test, performed with GraphPad Prism 9 software.

### Data Sharing Statement

For original data, please contact.

## Results

### Actin levels are reduced in *MYH9*-RD RBCs

To investigate the symptoms caused by *MYH9*-RD mutations, we collected whole blood samples from three *MYH9*-RD patient donors (R702S, D1424N and R1464C) and healthy human donors simultaneously. All three *MYH9*-RD patients exhibited significant enlargement of platelets in Wright-Giemsa stained smears (Figure S1) and had reduced platelet counts (Table 1). We analyzed the levels of NMIIA HC and cytoskeleton proteins in the whole blood of the patients using Western blot. Unexpectedly, we found that the actin levels in the whole blood of *MYH9*-RD patients were noticeably lower than those in the healthy controls. When equal amount of total proteins were loaded for the whole blood samples of the patients and the healthy controls, the actin levels in the whole blood samples of R702S, D1424N, and R1464C patients were approximately 50%, 69%, and 71%, respectively, of those in the controls (Figure 1A and 1B).

**Table 1.** Complete Blood Counts of *MYH9*-RD patients and the reference ranges for healthy individuals.

|  | R702S (♀) | D1424N (♂) | R1464C (♀) | Reference ranges |
| --- | --- | --- | --- | --- |
| PLT ($\times 10^9/L$) | 24 | 14 | 105 | 125-350 |
| RBC ($\times 10^{12}/L$) | 4.84 | 5.13 | 4.41 | 4.0-5.5 (♂), 3.8-5.1 (♀) |
| WBC ($\times 10^9/L$) | 4.63 | 3.7 | 9.41 | 3.5-9.5 |
| MCV (fl) | 82.8 | 92 | 88.4 | 82-100 |
| MCH (pg) | 27.1 | 30.5 | 30.4 | 27-32 |
| MCHC (g/L) | 328 | 332 | 344 | 320-360 |
| RDW (%) | 15.8 | 13.6 | 12.3 | 11-16 |
| HCT (%) | 40.1 | 47.2 | 39 | 35-50 |
| HGB (g/L) | 132 | 157 | 134 | 120-160 (♂), 110-150 (♀) |

Note: PLT, platelet count; RBC, red blood cell count; WBC, white blood cell count; MCV, mean corpuscular volume; MCH, mean corpuscular hemoglobin; MCHC, mean corpuscular hemoglobin concentration; RDW, red cell distribution width; HCT, hematocrit; HGB, hemoglobin. ♂, male; ♀, female.

**Figure 1.**
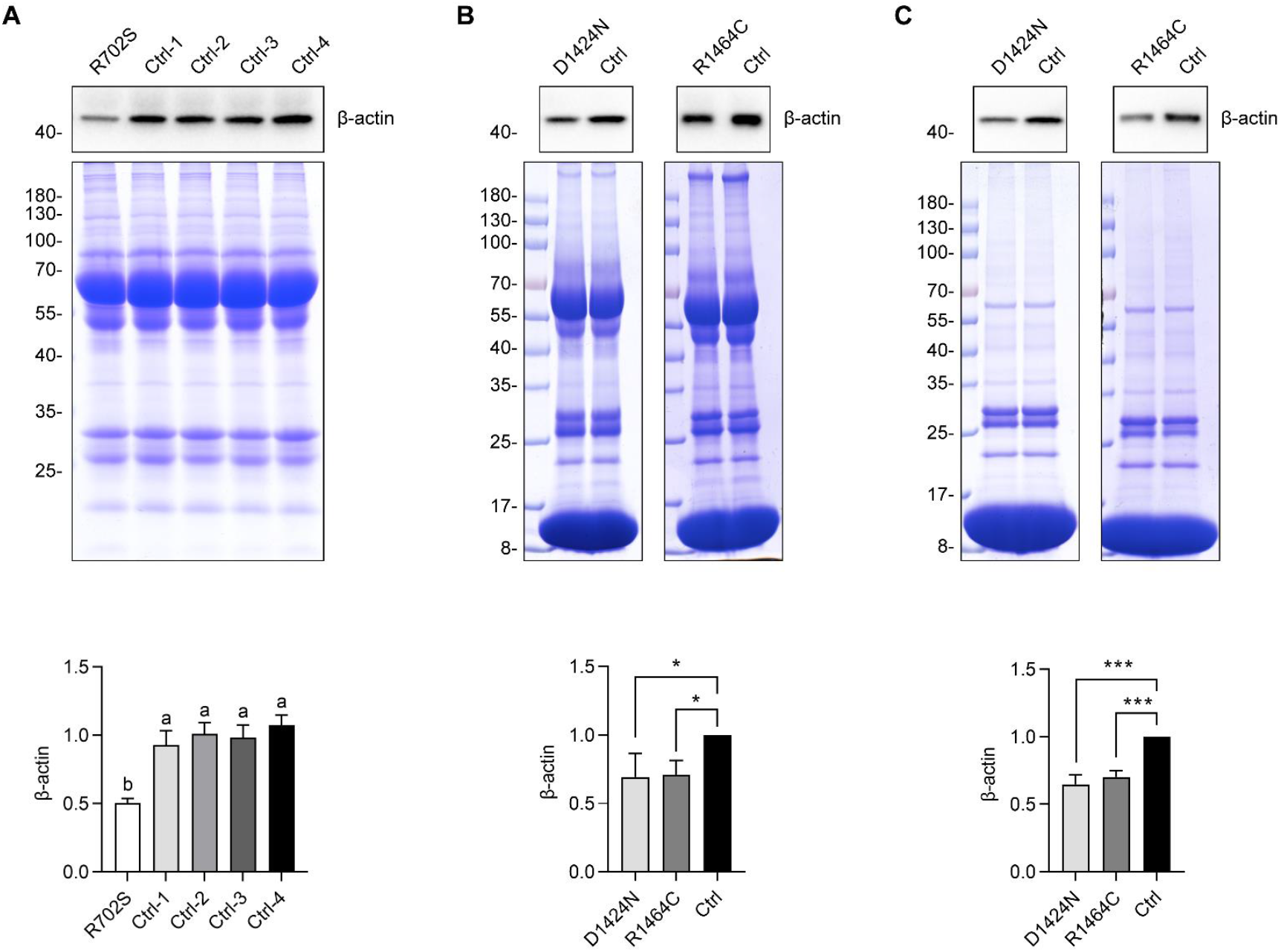
Actin levels in whole blood and RBCs of *MYH9*-RD patients. (A, B) Comparisons of actin levels in whole blood of *MYH9*-RD patients and healthy controls. Actin levels in whole blood were detected by Western blot using anti-actin antibody (top) and total proteins were detected by SDS-PAGE and Commassie blue staining (middle). Bottom shows quantification of actin levels in whole blood of *MYH9*-RD patients and healthy controls detected by Western blot. The results are the mean ± SD of three independent repeats. (A) The relative levels of actin of R702S patient and healthy controls were normalized to the average value from 4 healthy controls. The statistical differences among the samples were determined using one-way ANOVA with Bonferroni post hoc test (P < 0.05). (B) The relative actin levels of D1424N and R1464C patients were normalized to that of the corresponding healthy control. Data are the mean ± SD of three independent experiments with one-way ANOVA with Bonferroni post hoc test (*P < 0.05). (C) Comparisons of actin levels in RBCs of *MYH9*-RD patients and healthy controls. Actin levels in RBCs were detected by Western blot using anti-actin antibody (top) and total proteins were detected by SDS-PAGE and Coomassie blue staining (middle). Bottom shows quantification of actin levels in RBCs of *MYH9*-RD patients and healthy controls detected by Western blot. The relative actin levels of D1424N and R1464C patients were normalized to that of the corresponding healthy control. Data are the mean ± SD of three independent experiments with one-way ANOVA with Bonferroni post hoc test (***P < 0.001).

Given that RBCs are the most abundant cell type in blood, we speculated that the actin in whole blood is primarily derived from RBCs and the decrease in actin level observed in the whole blood of *MYH9*-RD patients is mainly due to RBCs. Indeed, Western blotting showed that ∼75% of actin in whole blood is derived from RBCs (Figure S2). We then isolated RBCs from the whole blood of D1424N and R1464C patients and analyzed the actin levels using Western blot. As expected, we found that the actin levels in both D1424N and R1464C RBCs were significantly lower than in the healthy controls (Figure 1C). Densitometry of the Western blot showed that the levels of actin in D1424N and R1464C RBCs were approximately 65% and 70% of that in the healthy human RBCs, respectively (Figure 1C).

Due to lack of isolated R702S RBC samples, we were unable to directly determine the actin level in R702S RBCs. However, we were able to estimate the actin levels in R702S RBCs based on the fact that RBCs contribute ∼75% actin in the whole blood of healthy control and the actin level in the R702S whole blood was reduced to 50% of the normal level. We expected that the actin levels in R702S RBCs were at most 67% of normal levels, even all actin in the R702S whole blood is derived from the RBCs. Therefore, we conclude that the actin levels are reduced in the RBCs of all three *MYH9*-RD patients, i.e., R702S, D1424N, and R1464C.

We also determined the NMIIA levels in the whole blood and the RBCs of *MYH9*-RD patients by Western blot. Compared with the healthy control, the NMIIA levels in the whole blood of *MYH9*-RD patients were significantly reduced, i.e., 58% for D1424N, 33% for R1464C, and 52% for R702S of those in healthy controls (Figure S3A). The NMIIA levels in the RBCs of D1424N, and R1464C were 60% and 36%, respectively, of those in healthy controls (Figure S3B). We were not able to estimate the NMIIA levels in R702S RBCs, because NMIIA in RBCs constitutes only a minor fraction of the total NMIIA in whole blood (Figure S2).

### The levels of several membrane skeleton proteins are reduced in *MYH9*-RD RBCs

In RBCs, actin exists in short filaments with a highly uniform length of ∼37 nm. Short actin filaments and several associated proteins assemble into junctional complexes interconnected by long (α1β1)_2_-spectrin tetramers, forming a two-dimensional cytoskeletal network beneath the plasma membrane of RBCs. Because the stoichiometric ratio of membrane skeletal proteins in RBCs is largely constant, we expected that a reduction of actin levels in RBCs would lead to a decrease in the levels of other membrane skeleton proteins.

To test this hypothesis, we examined the levels of several membrane skeleton proteins, including α-spectrin, tropomodulin, P4.1 and dematin, in the whole blood and RBCs of *MYH9*-RD patients using Western blot (Figure 2). In whole blood, α-spectrin levels in the three patients were approximately 75% of those in healthy controls and tropomodulin levels in R702S, D1424N, and R1464C were approximately 54%, 76%, and 78%, respectively, of those in healthy controls. P4.1 levels in D1424N and R1464C were similar to those in healthy controls, whereas that in R1464C was approximately 74% of those in healthy controls. The levels of dematin in three patients were similar to those of healthy controls. In RBCs, α-spectrin levels and tropomodulin levels in both D1424N and R1464C were reduced (76% in D1424N and 73% in R1464C for α-spectrin; ∼74% in both D1424N and R1464C for tropomodulin). The levels of P4.1 and dematin in both D1424N and R1464C were similar to those in health controls. Because α-spectrin and tropomodulin in whole blood are majorly derived from RBCs (Figure S2), we inferred that the reduction of those proteins in whole blood of R702S patients originates from RBCs.

**Figure 2.**
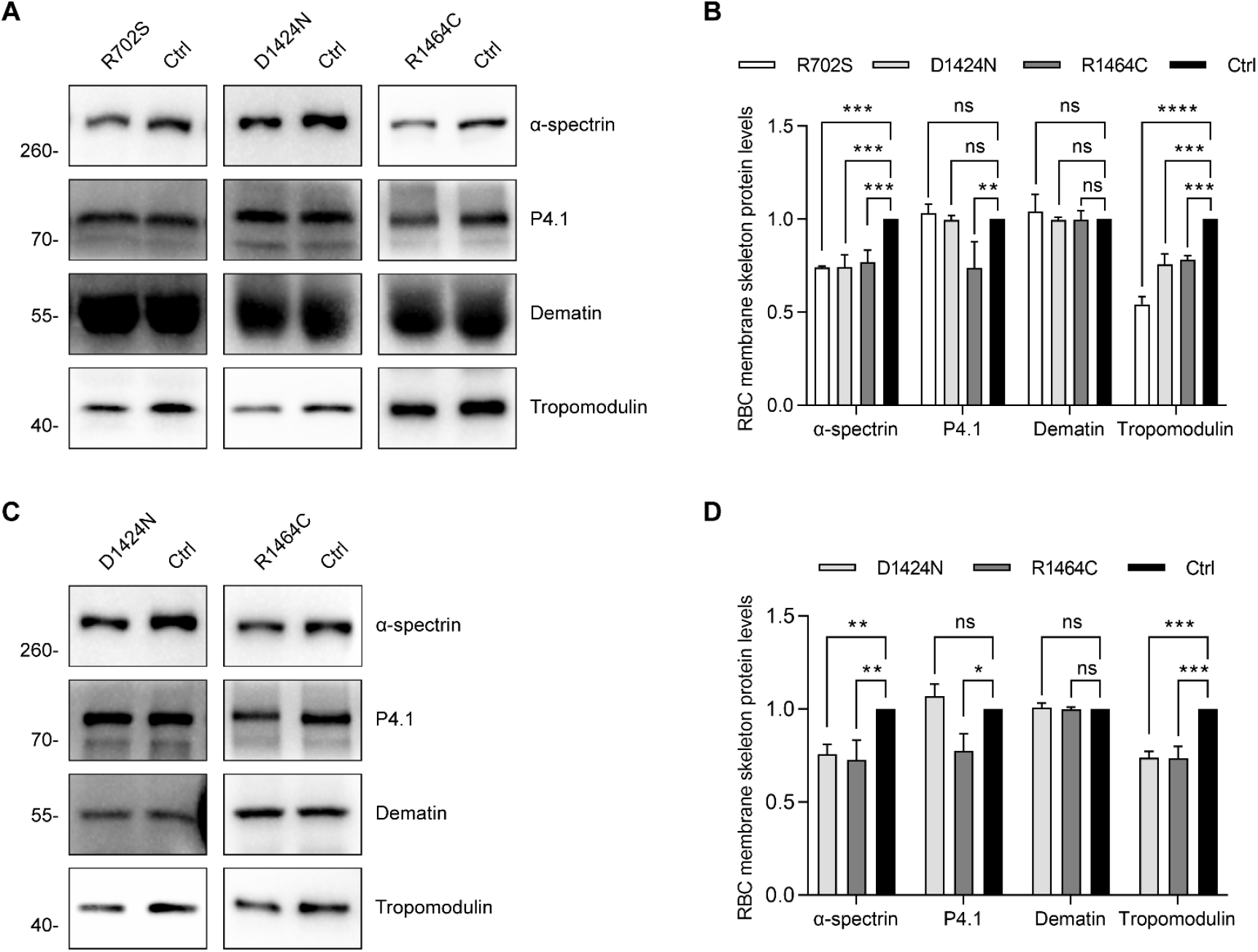
The membrane skeleton protein levels in whole blood and RBCs of *MYH9*-RD patients. (A) Comparison of the levels of RBC membrane skeleton proteins (α-spectrin, P4.1, dematin, and tropomodulin) in whole blood of *MYH9*-RD patients (R702S, D1424N, and R1464C) and healthy controls. Left, Western blot of RBC membrane skeleton proteins in whole blood samples of *MYH9*-RD patients and healthy controls. Right, relative amounts of RBC membrane skeleton proteins in the whole bloods. The levels of membrane skeleton proteins for patients are normalized to healthy controls. Data are the mean ± SD of 3 independent experiments with one-way ANOVA with Bonferroni post hoc test (ns, no significance, **P < 0.01, ***P < 0.001, ****P < 0.0001). The loaded proteins in SDS-PAGE with Coomassie blue staining are shown in Figure S4. (B) Comparison of the membrane skeleton protein levels in RBCs of *MYH9*-RD patients (D1424N and R1464C) and healthy controls. Left, Western blot of RBC membrane skeleton proteins in the RBCs of D1424N, R1464C, and healthy controls. Right, relative amounts of RBC membrane skeleton proteins in RBCs. The levels of membrane skeleton proteins for patients are normalized to healthy controls. Data are the mean ± SD of 3 independent experiments with one-way ANOVA with Bonferroni post hoc test (ns, no significance, *P < 0.05, **P < 0.01, ***P < 0.001).

### The membrane skeleton density in *MYH9*-RD RBCs is decreased

Given the reduced levels of several membrane skeleton proteins in *MYH9*-RD RBCs, we expected a lower density or abnormality of membrane skeleton networks in *MYH9*-RD RBCs. We used quick-frozen deep-etch and Cryo-SEM technique to examine the membrane skeleton networks in the RBCs of *MYH9*-RD and the healthy controls (Figure 3). The normal RBCs exhibit a relatively uniform network composed of filaments intersecting at multiple branching points, whereas the membrane skeleton lattice apertures in D1424N RBCs and R1464C RBCs were much larger and more variable than in normal subjects. Quantification of the lattice pore sizes revealed that the average lattice size in normal RBC membrane skeletons was approximately 2342 nm^2^, whereas the lattice pore sizes in D1424N and R1464C RBC membrane skeletons averaged approximately 4481 nm^2^ and 4984 nm^2^, respectively (Figure 3B).

**Figure 3.**
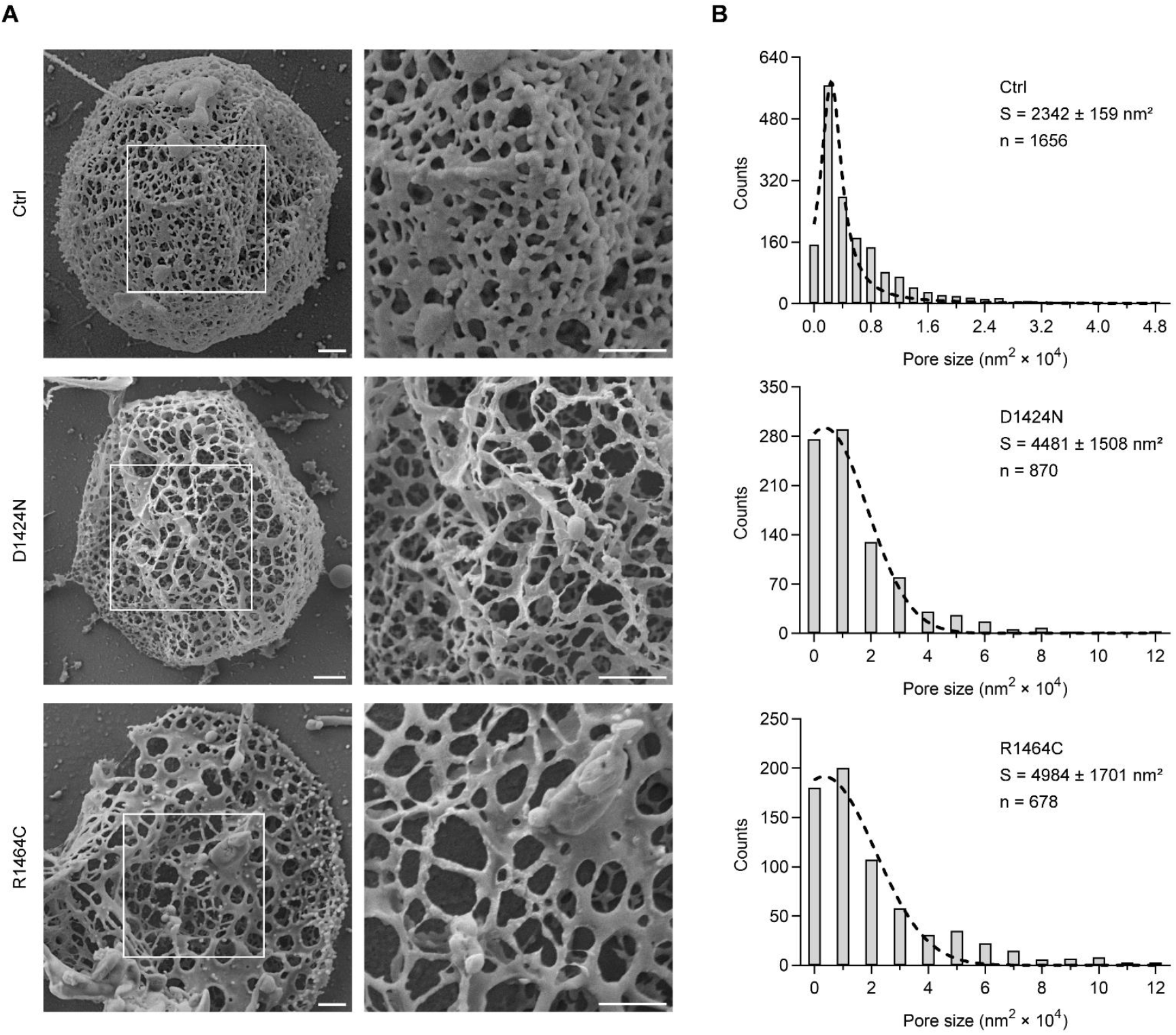
SEM images of RBC membrane skeleton of healthy control and *MYH9*-RD patients. (A) SEM images of the quick-frozen, deep-etched RBC membrane skeleton of healthy controls, D1424N, and R1464C. Scale bars = 500 nm. (B) Quantification of lattice pore sizes in the RBC membrane skeletons of healthy controls, D1424N, and R1464C. Only the pores located in the central 3/4 region of RBCs were quantified. Statistical data for each group are derived from 5 RBCs. “n” is the number of quantified pores.

### *MYH9*-RD RBCs produce greater deformation under hypertonic conditions compared to healthy controls

The biconcave disk morphology and deformation of RBCs rely on the membrane skeleton. Given the vast impact of the *MYH9*-RD mutations on the components and structure of RBC membrane skeleton, we expected some defects in the morphology and deformability of *MYH9*-RD RBCs. Wright-Giemsa-stained peripheral blood smears showed that approximately 20% of RBCs from both D1424N and R1464C patients exhibited abnormal shape with jagged edge, whereas very few RBCs from healthy controls did (Figure S5). We examined morphology of *MYH9*-RD RBCs and healthy controls under different osmotic conditions (Figure 4). Under hypotonic conditions with 0.675% NaCl, nearly all RBCs remained intact without undergoing osmotic lysis. Notably, abnormal crenated RBCs were seldom observed in both patients, suggesting the previously aberrant morphology may have been partially restored. Surprisingly, after isotonic treatment with 0.9% NaCl, crenated cells in both patients were also seldom observed, suggesting that the morphologically abnormal RBCs may have reverted to a normal shape. Under hypertonic conditions with 1.2% NaCl, both patients RBCs, as well as the healthy control, exhibited crenated morphology. Under hypertonic treatment with 1.5% NaCl, nearly all RBCs from the patients and the healthy control exhibited morphological changes. However, the deformation of the *MYH9*-RD RBCs was significantly more severe, with most showing significant damage. In contrast, healthy RBCs were relatively less damaged. Above observations suggest that the mechanical properties of *MYH9*-RD RBCs membrane are compromised, especially resistance to hypertonic pressure is reduced.

**Figure 4.**
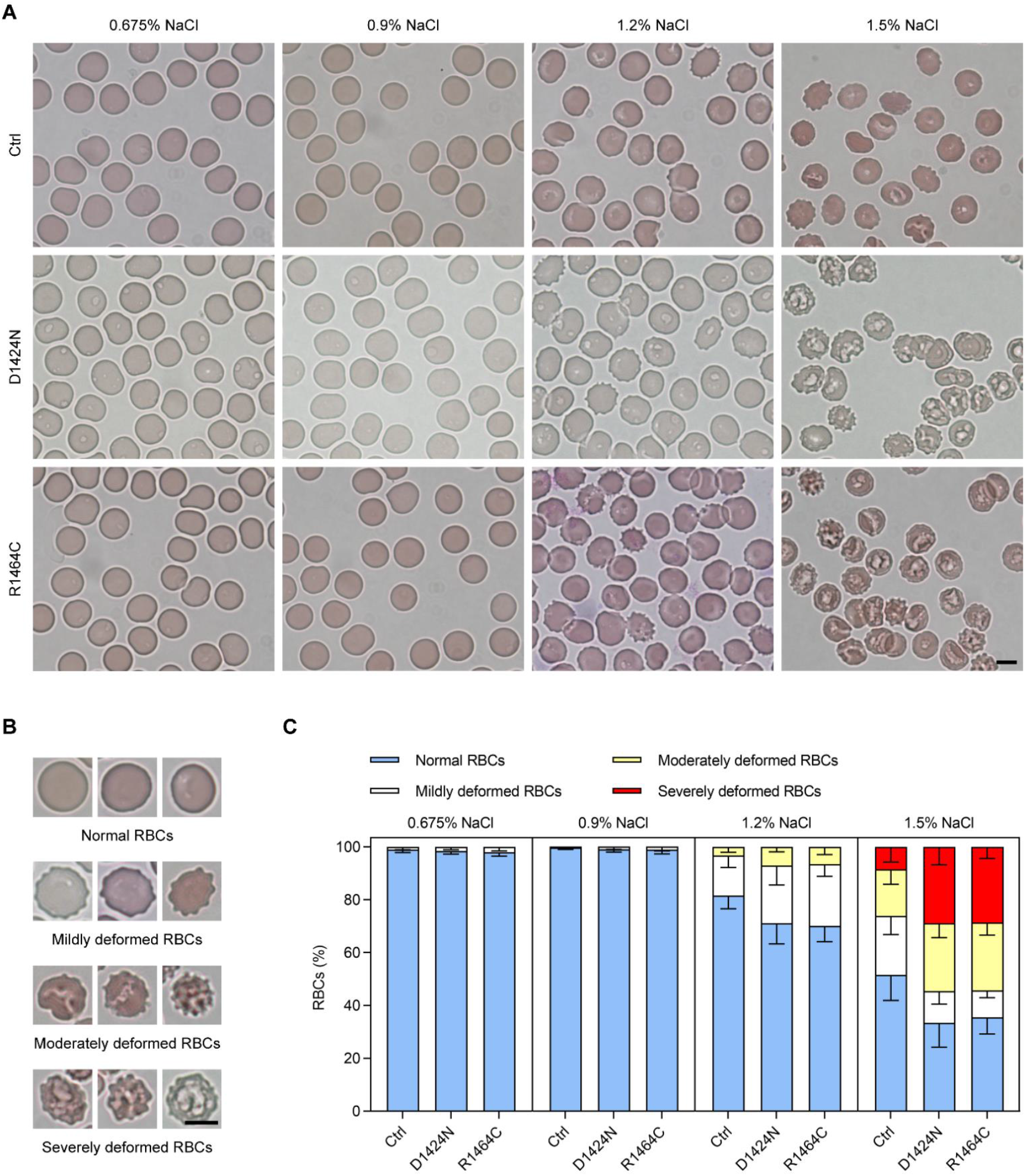
Wright-Giemsa-stained peripheral blood smears from *MYH9*-RD patients and normal subjects under different osmotic conditions. (A) Representative images of smears from a normal control, and *MYH9*-RD patients with D1424N or R1464C mutations under different NaCl concentration conditions. Scale bar = 5 μm. (B) Classification of RBC morphology based on the severity of deformation. (C) Quantification of RBC morphology under different NaCl concentration conditions.

## Discussion

Typical symptoms of *MYH9*-RD are macrothrombocytopenia and Döhle bodies in leukocytes ^1,6^. Here, we discovered a new symptom of *MYH9*-RD: decreased density in the membrane skeleton in *MYH9*-RD RBCs. We found that in all three *MYH9*-RD patients tested (R702S, D1424N, and R1464C), levels of at least three membrane skeleton proteins (actin, α-spectrin and tropomodulin) in RBCs are substantially decreased. We also found that *MYH9*-RD RBCs have enlarged lattice aperture of membrane skeleton network and abnormal deformability compared to healthy controls. We propose that defects in the membrane skeleton network in RBCs is a common symptom in *MYH9*-RD.

In this study, we were able to detect altered actin content in *MYH9*-RD RBCs because our Western blot results were normalized to the amount of total proteins rather than to actin level. β-actin is one of the most commonly used loading control during a Western blot, as it is expressed at high levels within most eukaryotic cell types and is generally unaffected by experimental conditions. However, this is not the case for *MYH9*-RD RBCs. As actin levels decreased in *MYH9*-RD RBCs, so did α-spectrin and tropomodulin, consistent with the fact that these two proteins interact closely with actin in the RBC membrane skeleton network. On the other hand, we did not observe a clear reduction of other membrane skeleton proteins such as P4.1 and dematin in *MYH9*-RD RBCs. We expect that those extra membrane skeleton proteins may bind abnormally to actin or diffuse in the cytosol.

A recent study also reported that *MYH9*-RD mutations cause the abnormal morphology of RBCs ^7^. However, in contrast to our findings, they found that *MYH9*-RD mutations did not affect the composition and organization of RBC membrane skeleton proteins, including actin and α1-spectrin. This discrepancy may be attributed to the different sample handling and quantification control methods used in the current study as compared to the previous one. Firstly, using SDS-PAGE combined with protein staining (Coomassie blue staining), their study found that RBC ghosts from *MYH9*-RD patients contained normal levels of major membrane skeleton components such as α1/β1-spectrin, protein 4.1R, and actin. In contrast, we used RBCs rather than RBC ghosts as samples to quantify the levels of membrane skeleton proteins in the *MYH9*-RD RBCs. The preparation of RBC ghosts is more tedious and may mask the difference in the levels of membrane skeleton proteins in the RBCs. Secondly, their total internal reflection microscopy (TIRFM) images of actin and α1-spectrin beneath the plasma membrane show no detectable differences between the normal controls and *MYH9*-RD RBCs, whereas our quick-frozen, deep-etch scan electron microscopy of RBCs reveal the significantly enlarged lattice aperture in *MYH9*-RD RBCs. The distance between the actin nodes of RBC membrane skeleton network was estimated to be 40 nm to 200 nm ^33,34^, which cannot be resolved by conventional TIRFM.

The membrane skeleton is crucial for maintaining RBC morphology and deformation, and many RBC membrane diseases lead to abnormal deformability and fragility ^31,35^. It is noteworthy that the moderate but substantial reduction of membrane skeleton proteins in *MYH9*-RD RBCs only slightly affects the morphology and deformability of RBCs. RBCs are remarkably flexible and deformable, suggesting that the membrane skeleton network of RBCs is very resilient and may tolerate small defects. Interestingly, the *MYH9*-RD RBCs show fairly normal morphology and slight deformation under normal and hypotonic conditions, but are relatively more sensitive to hypertonic conditions. The RBC membrane skeleton consists of a number of basic mesh-like units, with membrane filaments interconnected to form a meshwork located beneath the cell membrane. The overall structure of RBC membrane skeleton resembles a cage, providing excellent tensile strength but relatively poor compressive strength. From this point of view, it is not unexpected that a moderate reduction in the density of membrane skeleton in *MYH9*-RD RBCs only slightly affects tensile strength, but greatly impairs compressive strength.

How *MYH9*-RD mutations lead to the reduction of membrane skeleton proteins in RBCs is a question that deserves further investigation. RBCs develop from committed stem cells through a process called erythropoiesis, which involves the expulsion of the nucleus and other intracellular organelles, as well as most of cytoskeletal structures, and the assembly of the membrane skeleton. It is possible that NMIIA plays a role in the assembly of the RBC membrane skeleton and abnormal activities of NMIIA mutants in *MYH9*-RD RBCs cause the abnormal assembly of membrane skeleton network. *MYH9*-RD mutations affect platelets, leukocytes, and RBCs. All these blood cell types differentiate from common myeloid progenitor cells or more primitive hematopoietic stem cells ^36,37^. It has been reported that *MYH9*-RD mutations can destabilize the folded state of NMIIA, making it more prone to adopting an active state^19^. This suggests that *MYH9*-RD mutations may lead to increased activities of NMIIA, potentially resulting in the premature differentiation of RBCs before proper maturation. This could also explain how abnormal proplatelet formation contributes to the macrothrombocytopenia and the rescue effects of blebbistatin ^16,17,20^. Taken together, we hypothesize that the proper assembly of membrane skeleton in RBCs depends on the normal activity of NMIIA and abnormal activities of NMIIA mutants in *MYH9*-RD RBCs cause the abnormal assembly of membrane skeleton network.

More than 100 *MYH9*-RD mutations have been identified so far. Those mutations are located either at the motor domain or at the filament-forming rod/tail domain of NMIIA. Based on the current finding that the three tested *MYH9*-RD patients, one with mutation at the motor domain (R702S) and two with mutations at the tail domain (D1424N and R1464C), all displayed reduced levels of membrane skeleton proteins actin, α-spectrin and tropomodulin in RBCs, we expect that reduced levels of membrane skeleton proteins in RBCs is a common symptom of *MYH9*-RD. Further studies are required to investigate the membrane skeleton in the RBCs of other *MYH9*-RD variants and to understand the underlined mechanism.

## Contribution

Conceived the overall research and supervised the study, X.d.L.; designed experiments, S.S., J.P., X.d.L.; conducted experiments, S.S., J.P., N.Z.; interpreted results, S.S., J.H., X.d.L.; collected and managed patient material and data, X.H.J., T.N.Z., Z.J.K., J.H.; wrote the manuscript, S.S., X.d.L.; critically reviewed and edited the manuscript, S.S., N.Z., J.H., X.d.L..

## Acknowledgements

We are very grateful to the *MYH9*-RD patients who contributed the blood samples. This research was supported by Beijing Natural Science Foundation (7232104), National Natural Science Foundation of China (31970657), and S&T Program of Hebei (22377772D).

