## Supplementary materials for "The membrane skeleton density of red blood cells in *MYH9*-related disease patients is decreased"

#### Supplemental Data

##### Figure S1

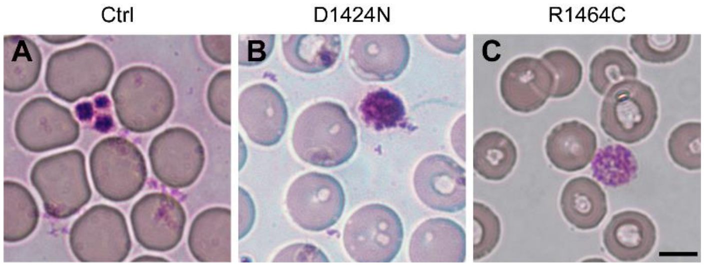


**Figure S1. Wright-Giemsa-stained peripheral blood smears from healthy control, D1424N, and R1464C.** The *MYH9*-RD patients exhibit significant enlargement of platelets. Scale bar = 5 μm.

##### Figure S2

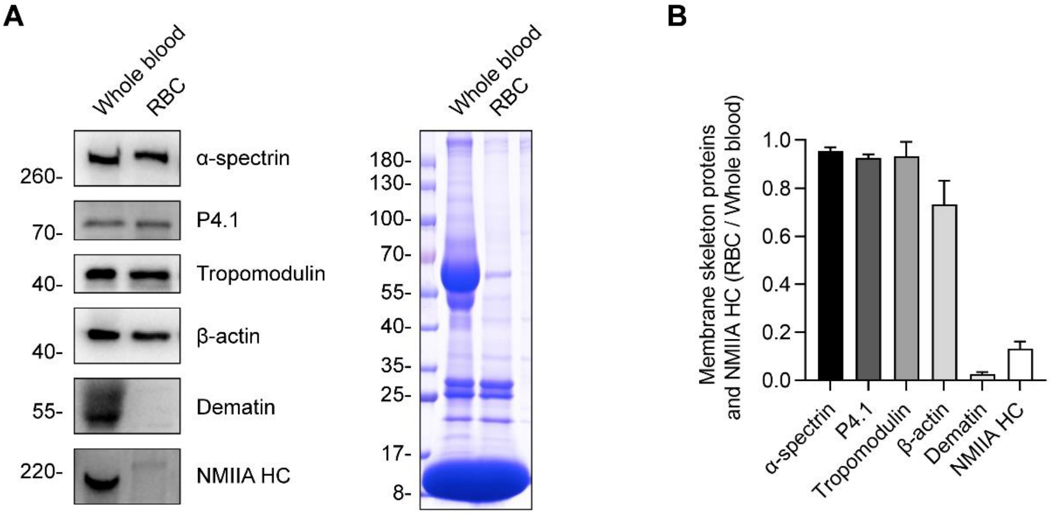


**Figure S2. Proportions of membrane skeleton proteins and NMIIA HC in RBCs relative to their levels in whole blood.** Analyses of the proportions of membrane skeleton proteins and NMIIA HC in RBCs relative to their total levels in whole blood, showing that with the exception of dematin and NMIIA HC, which are expressed at very low levels in RBCs, the remaining membrane skeleton proteins account for the vast majority of these proteins in whole blood. (A) Western blot (left panel) of membrane skeleton proteins and NMIIA HC in whole blood and in RBCs. Note that RBCs are isolated from whole blood, with the final volume adjusted to match that of the original blood sample. SDS-PAGE (right panel) of total proteins of whole blood and RBC samples used in left panel is served as loading control. Equal volumes are loaded for CBB staining. (B) Quantification of Western blot in (A). The ratios were calculated by dividing the band intensity in RBCs by the corresponding band intensity in whole blood. Data are the mean ± SD of three independent experiments.

### Figure S3

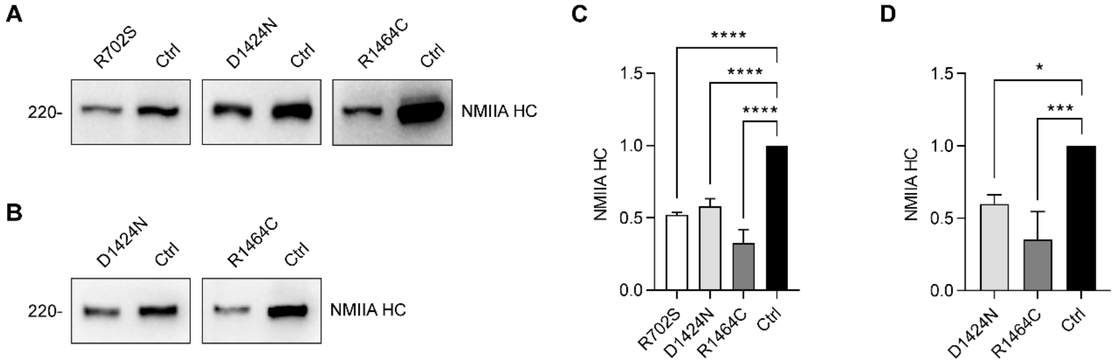


**Figure S3. The levels of NMIIA HC in *MYH9*-RD patients are reduced.** (A) Western blot of NMIIA HC in whole blood samples from *MYH9*-RD patients and healthy controls, showing NMIIA level is reduced in *MYH9*-RD whole blood compared to controls. (B) Western blot of NMIIA HC in RBCs from D1424N, R1464C and healthy controls, demonstrating NMIIA levels are reduced in D1424N and R1464C RBCs compared to healthy controls. (C) and (D) Quantification of western blot in (A) and (B), respectively. The levels of NMIIA HC for patients are normalized to the values from healthy controls. Data are the mean ± SD of 3 independent experiments with one-way ANOVA with Bonferroni post hoc test (*P < 0.05, ***P < 0.001, ****P < 0.0001).

### Figure S4

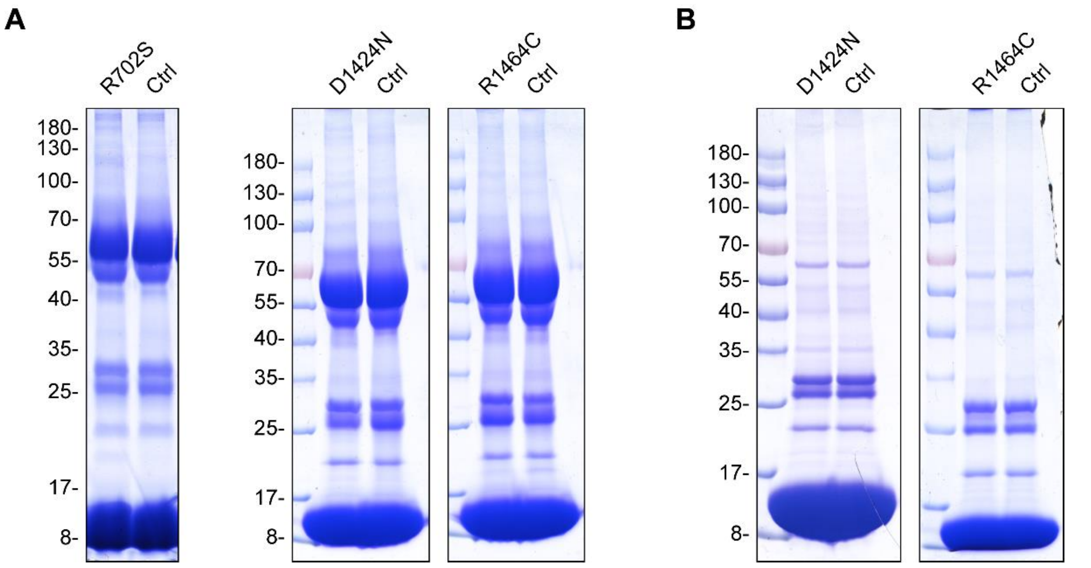


**Figure S4. Coomassie blue staining of the total proteins loaded in SDS-PAGE for Western blot in Figure 2 and Figure S3.** (A) Whole blood samples from R702S, D1424N, and R1464C, as well as the relevant controls. (B) RBC samples from D1424N and R1464C, as well as the relevant controls.

### Figure S5

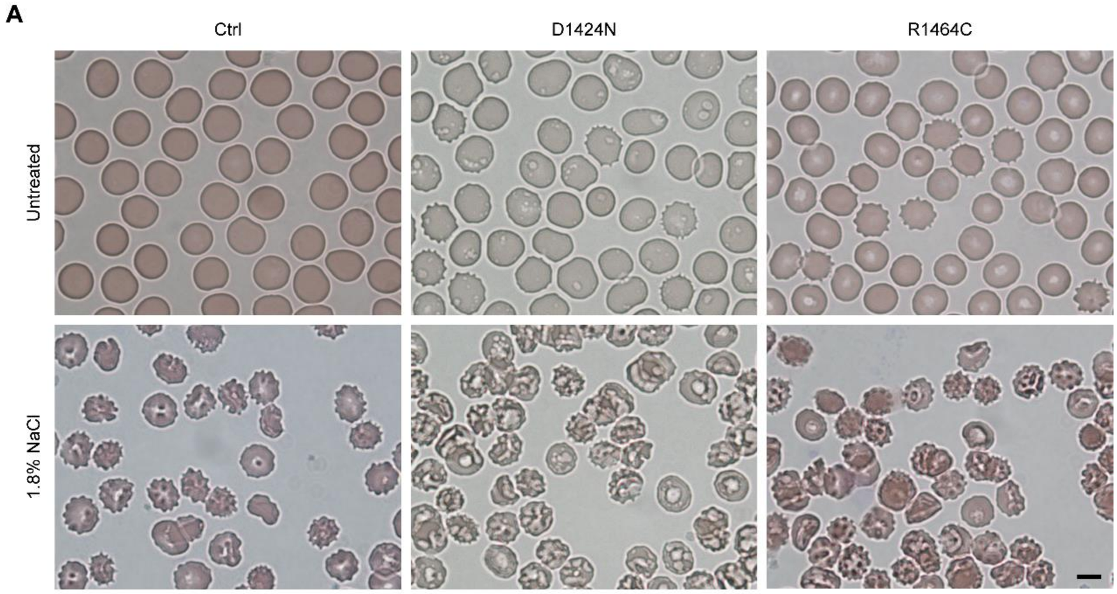


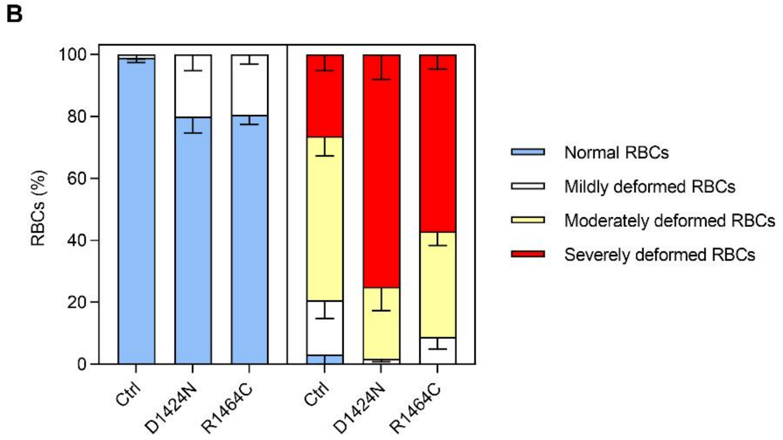


**Figure S5. Wright-Giemsa-stained peripheral blood smears from *MYH9*-RD patients and health control.** Wright-Giemsa-stained peripheral blood smears from D1424N and R1464C patients reveal RBCs with abnormal crenated morphology and varying deformability and fragility in response to hypertonic 1.8% NaCl conditions. (A) Representative images of peripheral blood smears from healthy controls, D1424N, and R1464C untreated (top) or treated with 1.8% NaCl (bottom). Scale bar = 5 µm. (B) Percentages of RBCs with varying degrees of deformation under untreated or 1.8% NaCl conditions.
